# Network mapping of root-microbe interactions in *Arabidopsis thaliana*

**DOI:** 10.1101/2020.11.24.397273

**Authors:** Xiaoqing He, Qi Zhang, Yi Jin, Libo Jiang, Rongling Wu

**Affiliations:** Beijing Advanced Innovation Center for Tree Breeding by Molecular Design, Beijing Forestry University, Beijing 100083, China; Center for Computational Biology, College of Biological Sciences and Technology, Beijing Forestry University, Beijing 100083, China; Center for Statistical Genetics, Departments of Public Health Sciences and Statistics, The Pennsylvania State University, Hershey, PA 17033, USA

**Keywords:** *Arabidopsis thaliana*, root microbe interaction, hub microbes, network mapping, QTL network, hub genes

## Abstract

Understanding how plants interact with their colonizing microbiota to determine plant phenotypes is a fundamental question in modern plant science. Existing approaches for genome-wide association studies (GWAS) are based on the association analysis between host genes and the abundance of individual microbes, failing to characterize the genetic architecture of microbial interactions that are thought to a determinant of microbiota structure, organization, and function. Here, we implement a behavioral model to quantify various patterns of microbe-microbe interactions, i.e., mutualism, antagonism, aggression, and altruism, and map host genes that modulate microbial networks constituted by these interaction types. We reanalyze a root-microbiome data involving 179 accessions of *Arabidopsis thaliana* and find that the four networks differ structurally in the pattern of bacterial-fungal interactions and microbiome complexity. We identify several fungus and bacterial hubs that play a central role in mediating microbial community assembly surrounding *A. thaliana* root systems. We detect 1142 significant host genetic variants throughout the plant genome and then implement Bayesian networks (BN) to reconstruct epistatic networks involving all significant SNPs and find 91 hub QTLs. Gene annotation shows that a number of the hub genes detected are biologically relevant, playing roles in plant growth and development, resilience against pathogens, root development, and improving resistance against abiotic stress conditions. The new model allows us to better understand the underlying mechanisms that govern the relationships between plants and their entire microbiota and harness soil microbes for plant production.

## INTRODUCTION

The microbiota has been widely thought to be an important determinant of various natural processes ranging from biogeographical cycling to human health. Many studies have characterized strong associations between the microbiota and a variety of human disorders (1–4), but research on how the microbiota impacts plant growth has not been conducted until recently (5, 6). Increasing evidence shows that the microbiota plays a pivotal role in promoting plants’ stress tolerance, determining plant productivity, improving the bioavailability of nutrients, and preventing invasion by bacterial pathogens (5, 7–12). Some bacteria can fix and preserve nitrogen in root nodules for plants(13, 14), whereas others even can modulate the timing of flowering of plants (15, 16) and save hosts from extinction (17). Under drought stress, root microbiomes can help crop plants maintain more sustainable production (10, 18).

While the microbiota affects the phenotypes of the hosts they colonize, the hosts can also shape the structure and function of the microbial communities (2, 4, 19, 20) in a way that is determined by environmental factors (2, 21). It has been commonly recognized that the microbiota and their hosts form complex but well-orchestrated interaction networks (22, 23). For example, the crop microbiome assembly is shaped by the host, even to a greater extent environmental factors (24). There is a greater variability among plant species or genotypes in their ability to recruit specific microbial communities (10, 25). Plant genes affect root metabolism, immune system functioning, and root exudate composition, which in turn influence the activity and structure of the root microbiome (26, 27). Recent studies provide a ‘cry-for-help’ hypothesis’ to explain that stressed plants assemble health-promoting soil microbiomes by changing their root exudation chemistry (28). It was still arguable that, to what extent, the overall microbiome is shaped by host genes.

Roots of healthy plants are colonized by multi-kingdom microbial consortia (22, 29, 30). The whole microbiome structure and function are determined by the pattern and strength of how the constituent microbes interact with each other through cooperation or competition (31). F. Getzke et al. (32) found that interactions between microbiota members, particularly bacterial-fungal interactions, contribute to plant health. Given that fungi have a strong influence on the structure of the root microbiome, characterizing both bacteria and fungi can enhance our understanding of the root microbiome (33). Several studies have identified highly interconnected ‘hub species’ in microbial networks that act as mediators between a host and its associated microbiome (2, 34). Yet, we are still unclear in which way microbes interact with each other within to shape polymicrobial communities (35). We know little about how microbiota members contribute to the establishment, stability, and resilience of microbial communities essential for the maintenance of plant health.

Understanding the fundamental questions described above requires integrated systems approaches (1, 13). Recently, with the application of next-generation sequencing, the microbiome data and host genetic data measured at unprecedented resolution have been increasingly available (30, 36–41). From these data, genome-wide association studies (GWAS) have been developed to systematically characterize the genetic underpinnings of microbiota-host associations in plants (33, 42, 43). However, traditional GWAS models can only detect the host QTLs responsible for the abundance of individual microbes, failing to disentangle the relationships of diverse microbial species and microbe-host interactions (23, 35, 44). In a previous study, we have integrated ecology theory and network science to derive a series of mathematical descriptors for measuring all possible types of microbe-microbe interactions, and quantify how each microbe interacts with every other through a web of cooperation and competition (45). These descriptors show their utility to to map the genetic architecture of the gut microbiota (45).

In this article, we report the application of our ecology-based network model to root-microbiota interactions in Arabidopsis. As a model system, Arabidopsis has been extensively studied, aimed to explore the interactions between microbial communities and hosts. In a GWAS including 179 accessions of *A. thaliana,* Bergelson et al (33, 42) identified associations between the abundance of individual microbes within root microbiomes and plant genotypes. By reanalyzing this dataset, we further reveal the intricate relationship among *A. thaliana* and its colonizing microorganisms. We identify hub microbes within the root microbiome, characterize how microbes interact across kingdoms, and illustrate to what extent those interactions affect plant health and how this process is governed by the host genes.

## RESULTS

### Co-occurrence networks of the root microbiota

We developed a behavioral model to define the strengths of mutualism, antagonism, aggression, and altruism between each pair of microbes, quantitatively described by *Z*_mu_, *Z*_an_, *Z*_ag_, and *Z*_al_, respectively (see Experimental Procedures). These mathematical descriptors have been biologically validated (45, 46), allowing us to reconstruct mutualism, antagonism, aggression, and altruism networks for the root microbiota of the *A. thaliana.* To reduce the complexity of the networks, we chose the most abundant 100 OTUs in bacteria and fungi, respectively, for the reconstruction of four types of bacterial-fungal networks (OTU1-100 are listed as bacteria and OTU100-200 as fungi). We calculated node-level topological properties (i.e., degree, betweenness, closeness and eigenvector) using the “igraph” R package. Bacterial and fungal co-occurrence network characteristics are listed in Table S1.

Interkingdom functional diversity among fungi and bacteria is important for maintaining ecosystem functioning (30) and microbial interkingdom interactions in roots can promote Arabidopsis survival (22). We calculated degree-centrality parameters to determine the relative importance of bacteria and fungi in each network. It indicates that bacteria are more central to the structure of the mutualism and altruism networks than fungi (Fig. 1A and 1D), as bacteria tend to have a higher number (i.e., degree) of network connections than fungi (P=0.00038; P=0.0001029). In contrast, fungi in the antagonism and aggression networks appears to have a higher number of network connections than bacteria (Fig. 1B and 1C; P = 0.0002128; P= 0.01961). We also calculated and compared interkingdom microbial OTU relationships (the number of links; edges information) among bacterial and fungal taxonomic groups in four interaction networks (Fig.1; Table S2). Bacterial OTUs belonging to classes Betaproteobacteria, Flavobacteriia, Actinobacteria, Gammaproteobacteria, and Alphaproteobacteria displayed a strong mutualistic relationship with fungal OTUs belonging to classes Leotiomycetes, Dothideomycetes, Sordariomycetes, Agaricomycetes, and others, respectively (Fig. 1A). Bacterial classes, such as Actinobacteria, Alphaproteobacteria, and Deltaproteobacteria and fungal classes, such as Leotiomycetes, Dothideomycetes and Sordariomycetes were antagonistic to each other (Fig. 1B). In the aggression network, there were three bacterial classes (Betaproteobacteria, Actinobacteria, and Flavobacteriia) which were aggressive to fungal classes (Leotiomycetes, Sordariomycetes, Dothideomycetes, etc.) (Fig. 1C). Bacterial classes including Actinobacteria, Alphaproteobacteria, Betaproteobacteria, and Gammaproteobacteria were altruistic to fungal classes, Dothideomycetes, Leotiomycetes, Sordariomycetes, Mortierellomycetes, etc. (Fig. 1D).

**Figure 1.**
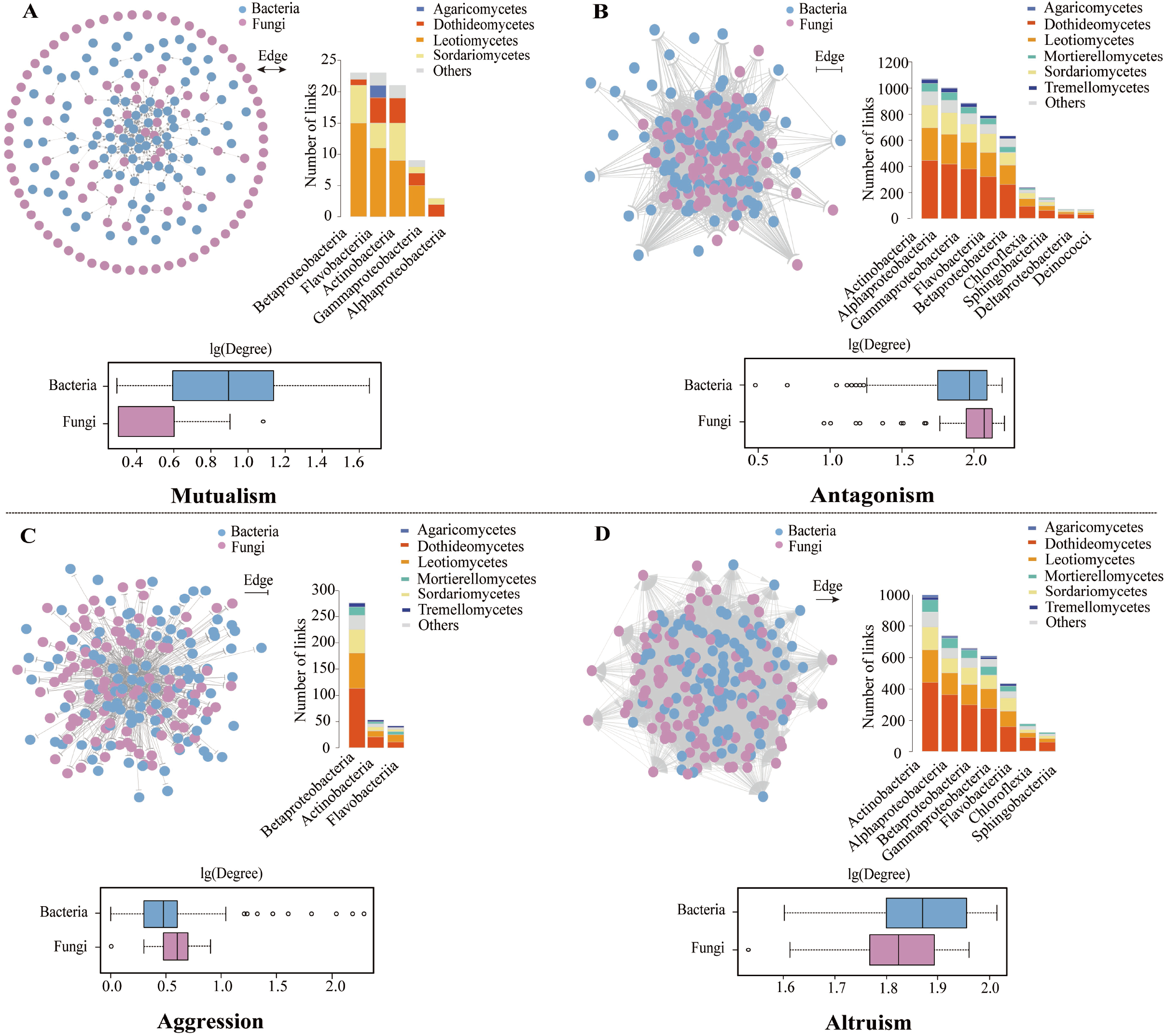
Microbial Z_mu_-based mutualism network (**A**), Z_an_-based antagonism network (**B**), Z_ag_-based aggression network (**C**), and Zal-based altruism network (**D**) at the OTU level of the root microbiome in *Arabidopsis thaliana*. In each network, bacteria and fungi are distinguished by different colors. The network analysis was performed in the “igraph” R package and visualized in Cytoscape v3.7.1. The number of links between root inter-kingdom microbes was given at the right. Bacterial and fungal OTUs were grouped at the class level and sorted according to number of edges between bacteria and fungi within each network. In boxplots, the fungal and bacterial degrees were calculated to determine the relative importance of bacteria and fungi in each network.

### Hubs of the co-occurrence network identification

Hub microbes are important in shaping microbial communities due to their critical roles in maintaining network function (2). The four networks differed structurally in the pattern of social links and the number of hub microbes. Fungal and bacterial OTUs that display the highest degree and the highest closeness centrality scores may serve as hub taxa to drive fungal-bacterial interaction equilibrium in *A. thaliana* roots (24) (Fig. 2A; Table S1).

**Figure 2.**
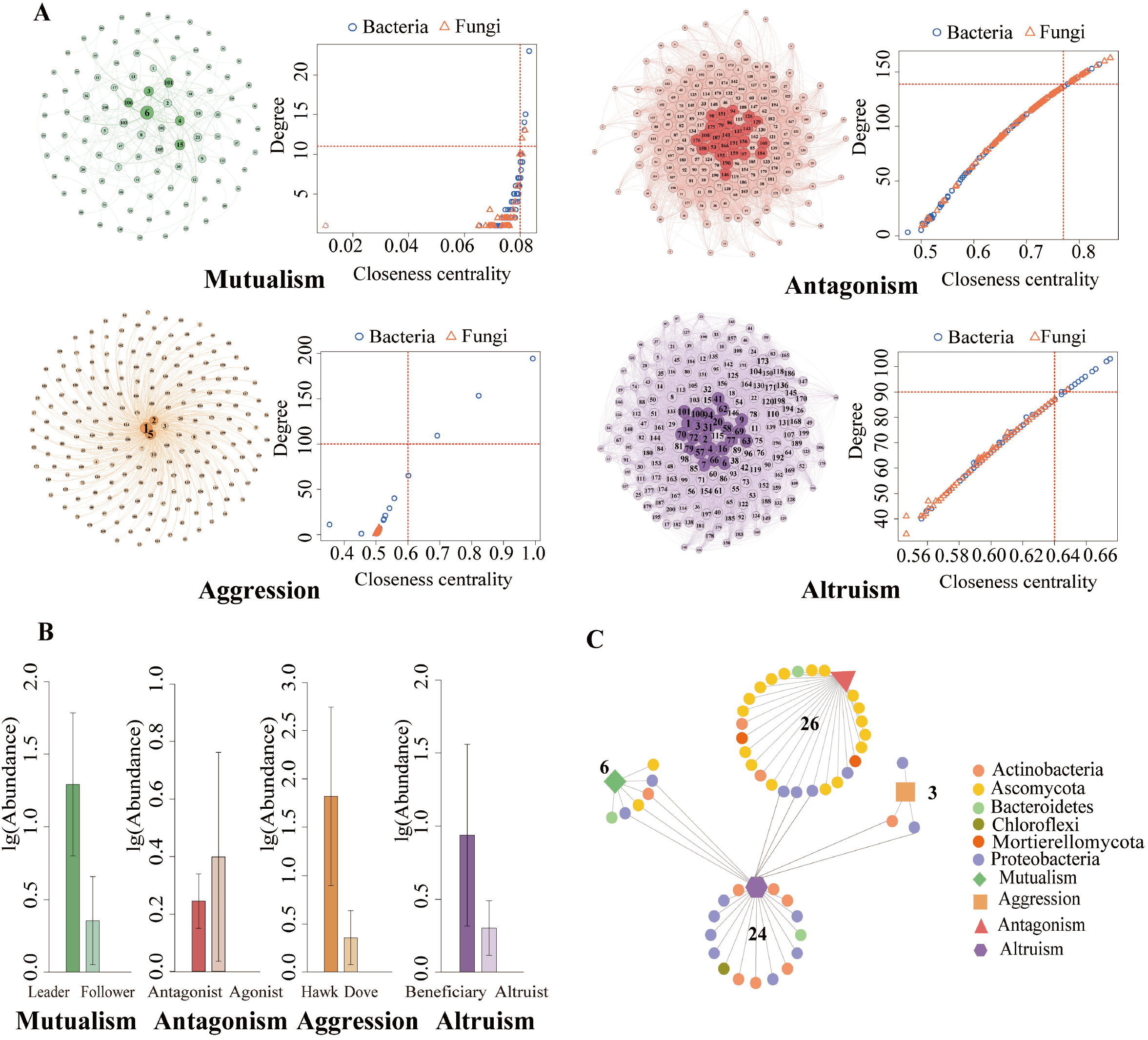
(**A)** The identity of each OTU is labeled by a number, 1 to 100 for bacteria and 101 to 200 for fungi. In each network, hub microbes are highlighted in dark circles. The distribution of ‘Hub microbes’ in four different microbial networks were based on degree and closeness centrality values. These two values of each OTU within each network were given at the right. The red dotted line represents the screening cutoffs of ‘Hub microbes’ corresponding to each network. (**B)** The abundance of hub microbes within each network. (**C)** The shared hub microbes within each microbial network. This ‘ shared network’ was represented in Cytoscape. Circle shapes represented hub microbes from each microbial network and irregular shapes represented different microbial network types. The edges were connected to hub microbes and microbial network. The distribution of ‘Hub microbes’ in four different microbial networks were based on degree and closeness centrality values. The red dotted line represents the screening cutoffs of ‘Hub microbes’ corresponding to each network. Visualization were done with Gephi for four microbial networks.

We identified six hub microbes, four bacteria and two fungi (nodes with degree >11 and closeness centrality values >0.08 in the network; P<0.01), which dominate the mutualism network. The four hub bacteria are classes Betaproteobacteria (2 OTUs), Sphingobacteriia, and Actinobacteria, and the two fungal hubs are phylum Ascomycota (2 OTUs). In the antagonism network, 26 hub microbes were found to act as ‘public enemies’, which were antagonistic to many more microbes than other microbes (degree >139 and closeness centrality values >0.78). The ‘agonists’ were observed to be more abundant than the ‘antagonists’ that are more combative (P<0.01; Fig.2B). In the aggression network, three OTUs belonging to the bacterial class Betaproteobacteria (2 OTUs) and Actinobacteria might represent the hub taxa (degree >100 and closeness centrality values >0.60; P<0.5). The hawks which are considered to aggressively repress others are abundant than the doves (those inhibited by others). The altruism network includes some ‘altruists’ (24 hub microbes; Fig. 2A; Table S1) that sacrificed their own growth by providing resources to beneficiaries (degree >90 and closeness centrality values >0.64). The hub microbes (altruists) are less abundant than the beneficiaries (Fig. 2B; P<0.01).

A total of 59 OTUs were identified as hub species, which were mainly from bacterial phyla proteobacteria (21 OTUs), Actinobacteria (12 OTUs), Chloroflexi (1 OTU), Bacteroidetes (3 OTUs) and fugal phyla Ascomycota (20 OTUs), Mortierellomycota (2 OTUs) (Table S1). The altruism network shares 4, 3, and 2 phyla with the mutualism, antagonism, and aggression networks, respectively (Fig. 2C).

### Mapping Root-microbe interactions

We calculated six centrality indices including connectivity (Con), closeness (C(u)), betweenness (B(u)), eccentricity (E(u)), eigencentrality (G(u)), and PageRank (P(u)) (Fig. 3) using the formulas given in Jiang et al. (45). In the same network type, these indices exhibit differences among hosts and, also, the same index varies among network types. By treating each network index as a phenotype, we performed association mapping for the interaction networks (Fig. S1). Our model identified 1142 significant host genetic variants throughout the plant genome, which are responsible for centrality indices of each network, including 225 acting through mutualism, 845 through antagonism, 49 through aggression, and 23 through altruism (19.70% for mutualism, 73.99% for antagonism, 4.29% for aggression, and 2.01% for altruism) (Table S3). It appears that more variants control mutualism and antagonism than aggression and altruism.

**Figure 3.**
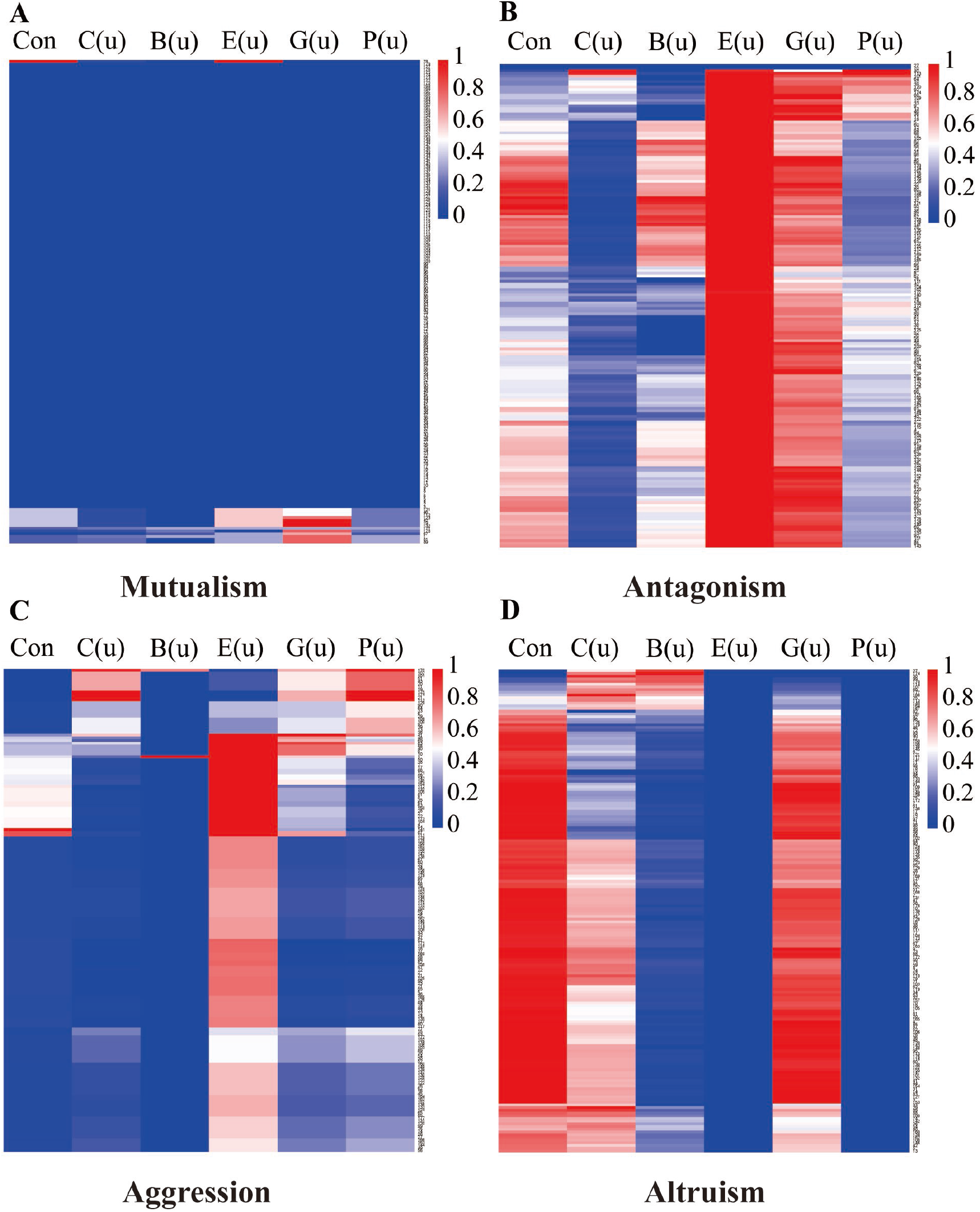
Heatmap of six emergent property indices constituting (**A**) Mutualism, (**B**) Antagonism, (C) Aggression, and (**D**) Altruism among 200 OTUs for network properties for each host.

### QTL networks

We implemented Jiang et al.’s (46) procedure to reconstruct Bayesian QTL networks among the significant SNPs detected to affect each type of microbial network and identified 91 hub QTLs (Fig. 4; Table 1). Through QTL network analysis, we can better characterize how a QTL mediates microbial cooperation or competition through its epistatic interactions with other QTLs. In the QTL network for the microbial mutualism network, we identify a hub QTL that affects connectivity QTL (Fig. 4A), annotated to gene *TMK3 (AT2G01820)* that orchestrates plant growth by regulation of both cell expansion and cell proliferation and as a component of auxin signaling (47). A hub QTL for the betweenness of microbial mutualism network is located in gene *IBR1(AT4G05530),* encoding indole-3-butyric acid response 1(IBR1). IBR1 are involved in root hair elongation (48). A hub QTL for the PageRank of the mutualism network represents gene *NTRB (AT4G35460)* encoding NADPH-dependent thioredoxin reductase. Thioredoxin is a key regulator of intracellular redox status that determine plant development in response to biotic and abiotic stress. Thioredoxin reductase (ntra ntrb) mutant alters both auxin transport and metabolism, causing a loss of apical dominance and reduced secondary root production, etc., largely regulated by auxin (49).

**Figure 4.**
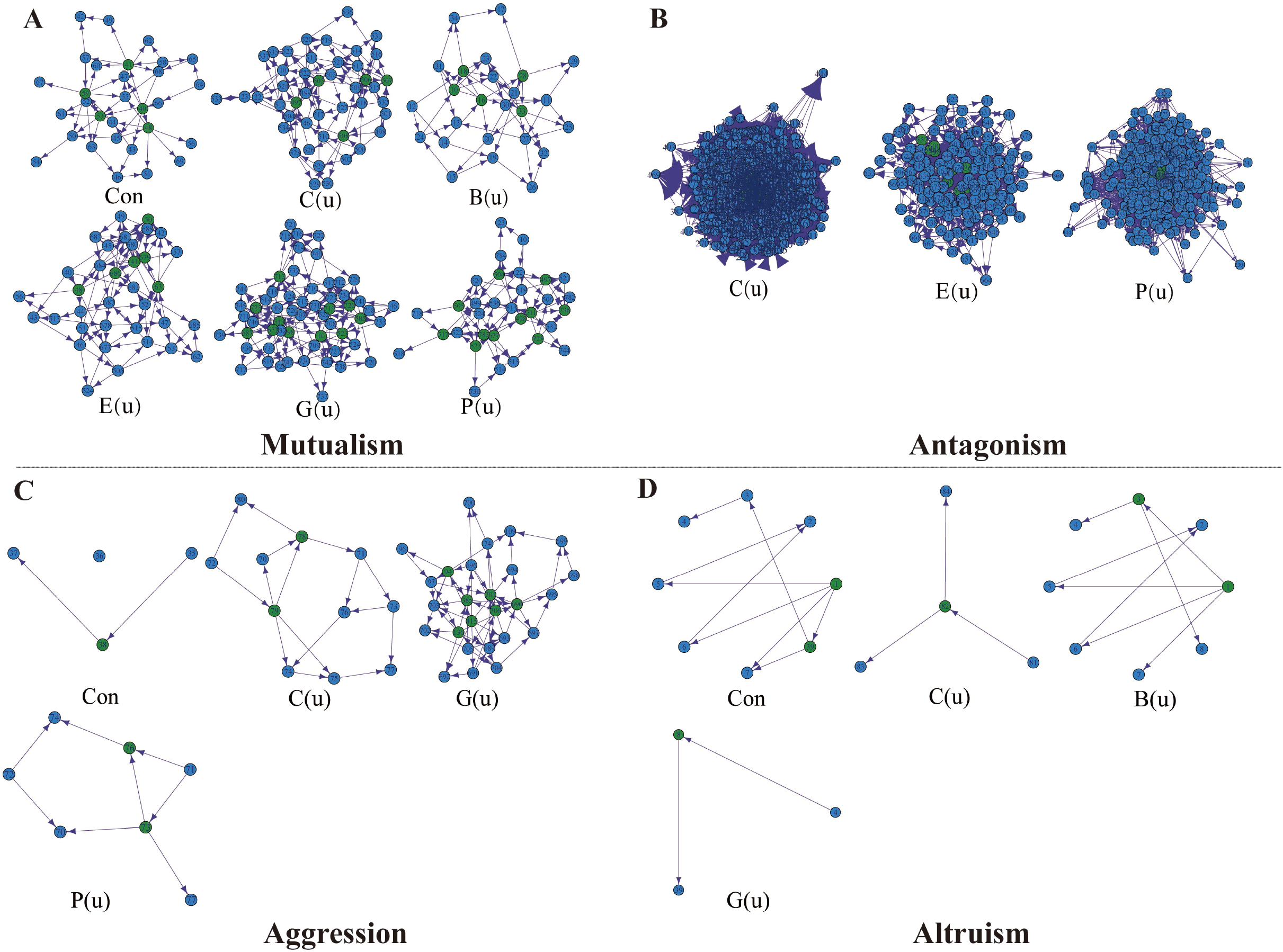
Bayesian QTL networks of significant SNPs mediating interaction networks of microbial mutualism (**A**), antagonism (**B**), aggression (**C**), and altruism (**D**). Hub QTLs within each genetic network are highlighted in green circles. The emergent properties of each microbial network are described by connectivity (Con), closeness (C(u)), betweenness (B(u)), eccentricity (E(u)), eigencentrality (G(u)), and PageRank (P(u)).

**Table 1.**
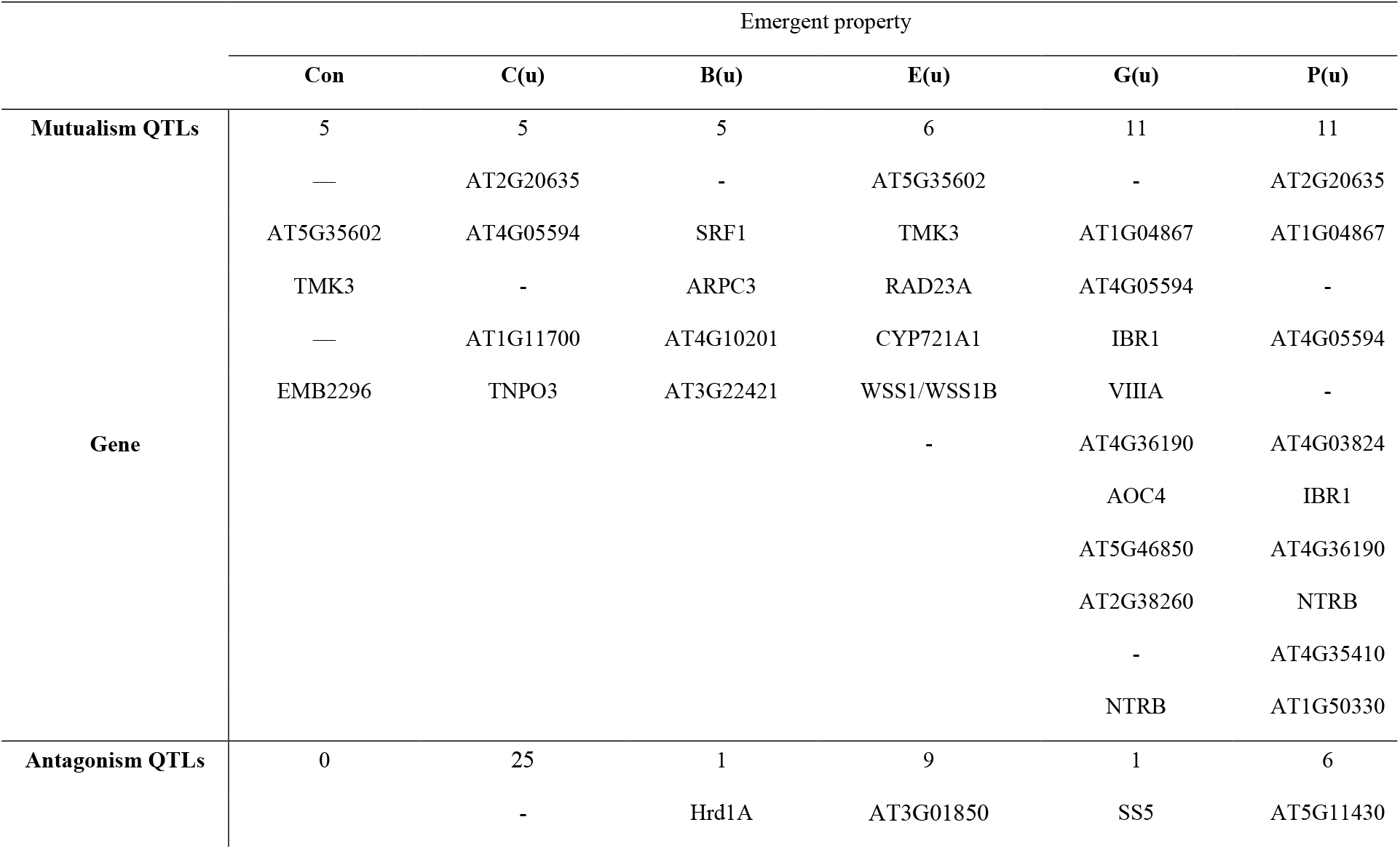
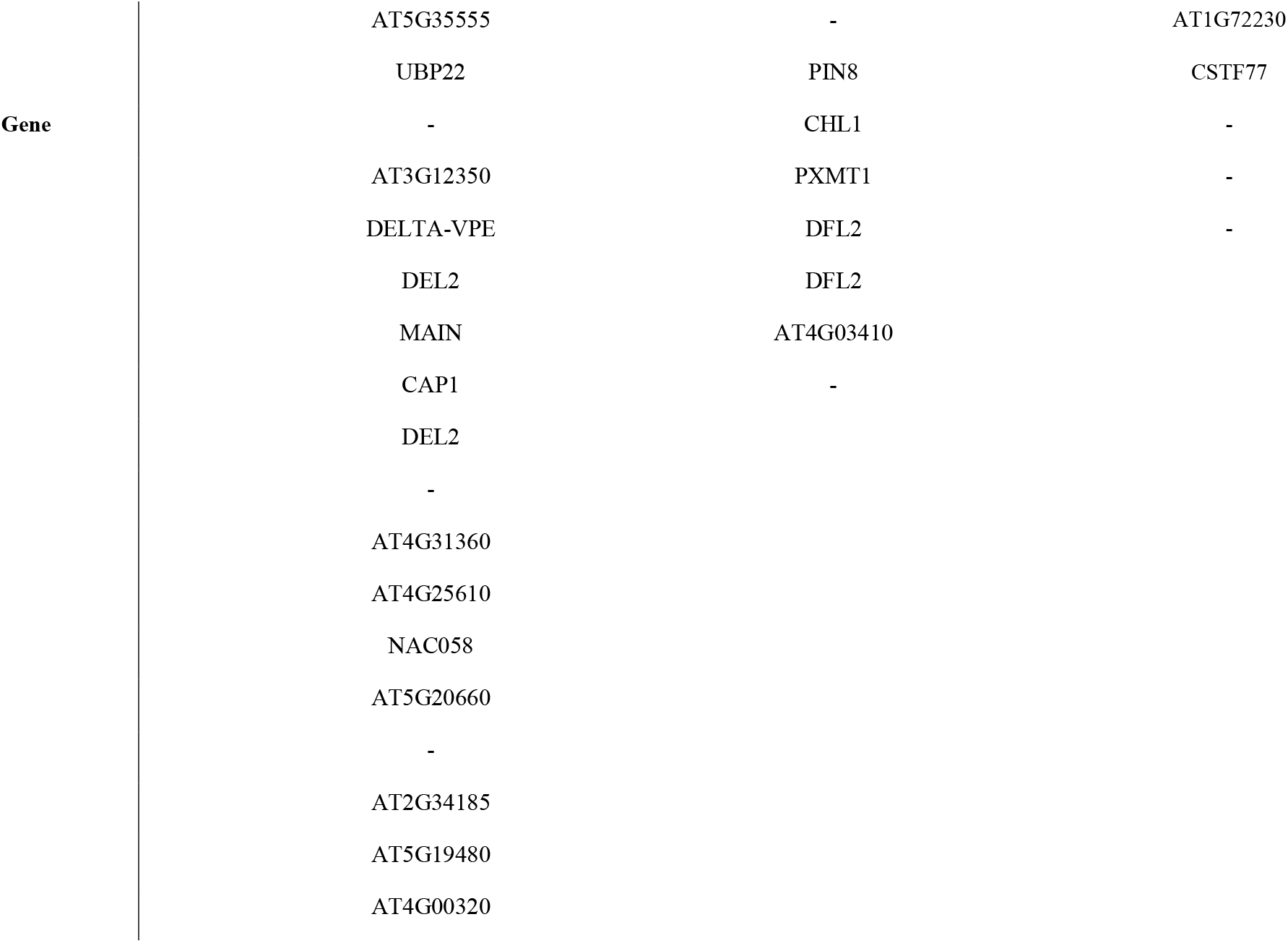
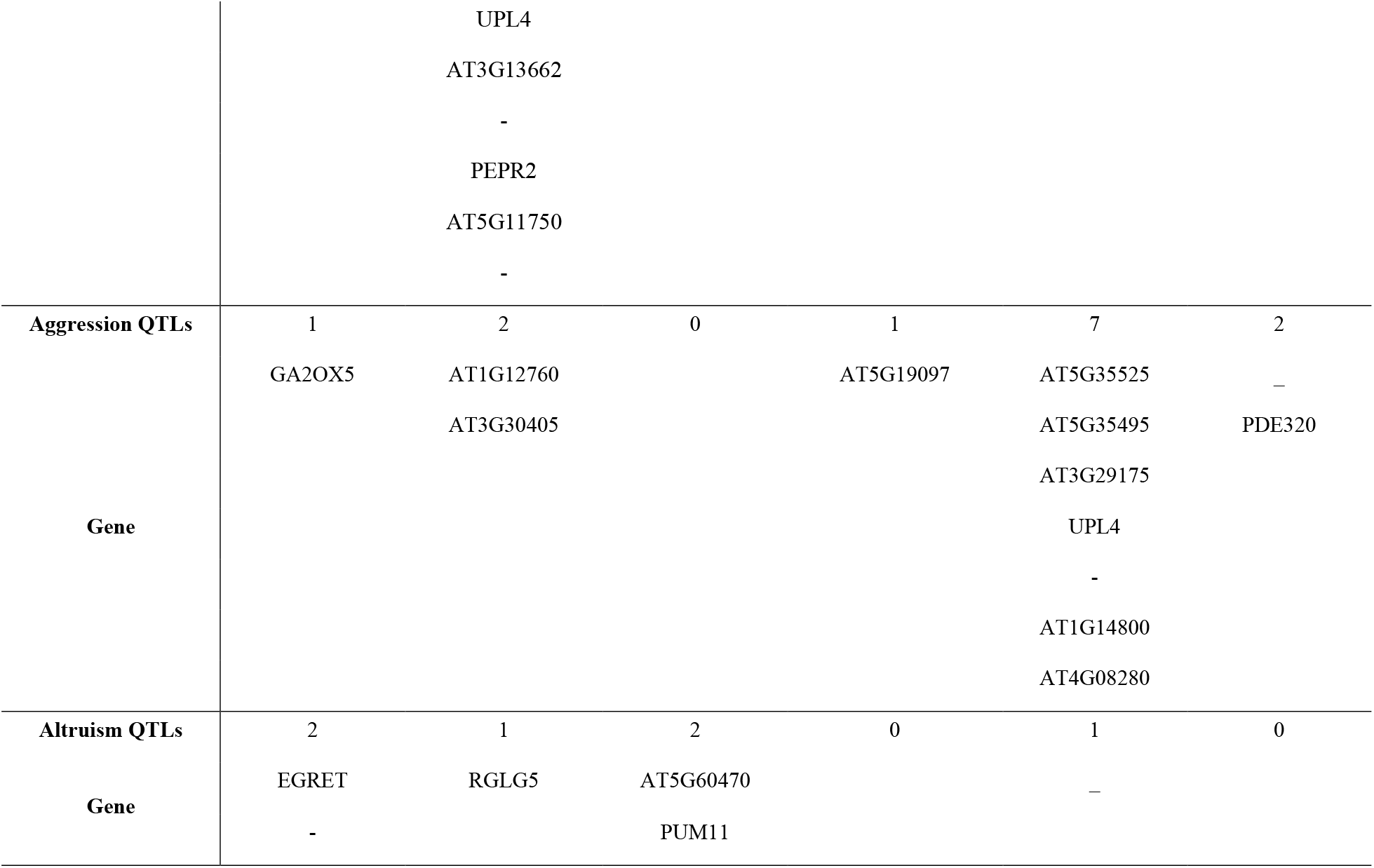
Numbers of mutualism, antagonism, aggression, and altruism QTLs that affect the emergent properties of ecological networks. The names of genes to which QTLs are annotated are given below.

In the QTL network for the microbial antagonism network, a closeness hub QTL acts like gene *UBP22(AT5G10790). UBP22* encodes a ubiquitin-specific protease, which plays roles in regulating plant development and stress responses (50). A hub QTL for the betweenness of the microbial antagonism network is located in gene *Hrd1A* which may be an important regulator of heat stress response in Arabidopsis (51). A hub QTL for the closeness of the microbial antagonism network acts like gene *PEPR2(AT1G17750)* encoding PEP1 receptor 2, which is transcriptionally induced by wounding and pathogen-associated molecular patterns and contributes to defense responses in Arabidopsis (52). *CHL1(AT5G40090)* is the hub QTL for the eigenvector of the antagonism network, which encodes disease resistance protein (TIR-NBS class). TIR-NBS protein is involved in disease resistance in Arabidopsis (53).

There are 12 pleiotropic genes including *UPL4(AT5G02880)*, which are detected to influence multiple types of microbial networks or properties (Table 1). *UPL4* encodes a ubiquitin-protein ligase, function additively in the regulation of plant growth and development, and positively modulate immune hormone salicylic acid (SA)-mediated basal and induced resistance responses (54).

## DISCUSSION

Plant rhizosphere is considered as the second genome of plants. As such, research on rhizosphere interactions has now become one of the hottest topics in modern biology and agriculture (55). However, the intrinsic principles governing the assembly of the root microbial community remain unclear (56). The potentially beneficial bacteria and fungi may serve as a valuable foundation for bio-fertilizer development in agriculture and forestry. Our understanding of the underlying mechanisms that govern the relationships between plants and their entire microbiota will be crucial for the benefit of sustainable plant growth (57).

### Co-occurrence network

Currently, network analysis has emerged as an extremely promising approach for modeling complex biological systems and can potentially provide deep and unique perspectives on microbial interactions and ecological assembly rules beyond those of simple richness and composition (58, 59). Properties of co-occurrence networks can reveal the intrinsic mechanisms of microbial interactions in response to environmental disturbance (38, 60). The connection and strength of the network even are crucial for the resistance to the pathogens. In this study, based on four descriptors, we successfully reconstructed corresponding 200-node networks accordingly. Each described root microbiome interactions according to a different ecological interaction metric. The four networks differ structurally in the pattern of bacterial-fungal interactions and microbiome complexity and the number of hub microbes.

### Microbial interkingdom interactions

In our study, bacteria have a higher degree in the mutualism and altruism networks and are more central to the structure of networks. Fungi have more connections in the antagonism and aggression networks, however, based on the directions of edges, bacteria still are more antagonistic and aggressive to fungi (Table S2). Understanding the interaction among different species within a community is one of the central goals in ecology (61). Microbial interkingdom interactions in roots could promote Arabidopsis survival. Bacterial communities aid in maintaining the microbial balance and protect host plants against the detrimental effects of filamentous eukaryotic microbes (22). A previous research showed that microbes tended to be positively related within kingdoms but negatively related between kingdoms (2). Besides bacteria and fungi, rhizosphere bacteriophages and protists also play roles in plant health (62, 63), which should be included in further research.

### Hub microbes

The identification of network hubs and their importance in microbial community structure has crucial implications for studying microbe–microbe interactions and can facilitate the design of strategies for future targeted biocontrol (2). Hub microorganisms have a regulatory influence on the network of microbial interactions, which can exert strong effects on microbiome assembly and serve as mediators between the plant and microbiome (2). According to centrality measurements, such as degree, closeness centrality and betweenness centrality, hub microorganisms are tightly connected within a co-occurrence network of (34). We identified hub microbes in four types of networks, mutualism, antagonism, aggression, and altruism. The most dominant taxa as hub microbes belong to bacterial phyla Proteobacteria (21 OTUs), Actinobacteria (12 OTUs) and fugal phyla Ascomycota (20 OTUs). This is consistent with the finding that Proteobacteria, Actinobacteria and Ascomycota are the most abundant phyla in plants and soil (24, 64). Actinobacteria is one of the bacteria whose dysbiosis in abundance in tomato rhizosphere causes the incidence of bacterial wilt disease (12).

### Host QTLs

Plant phenotypes are inextricably shaped by their interactions with microbes (1). In a well-designed GWAS study, J. Bergelson et al. (33) found a few significant QTLs that are associated with root microbial species richness and community structure, which are involved in plant immunity, cell-wall integrity, root and root-hair development. In this study, we used a newly developed network mapping model (45) to reanalyze Bergelson et al.’s (33) data, characterizing previously undetected QTLs that mediate microbial interactions. We found that most of the QTLs detected by the new model can be annotated to candidate genes with known biological functions including plant growth and development, resilience against pathogens, root development, and improved resistance against abiotic stress conditions (Table 1; Table S4). Understanding how microbes improve plant stress resistance will enhance our understanding of how plants survive in stress conditions. In the near future, it will be crucial to unravel the complex network of genetic, microbial, and metabolic interactions, including the signaling events mediating microbe-host interactions.

### Future prospects

Scientists has also linked the phyllosphere microbiome to plant Health (65) and found host genes could affect bacterial communities in the phyllosphere (20). Building synthetic microbiomes in plants has been proved to be useful for future research on plant-microbe interactions (66–71). Studies have showed the potential of microbiome adjustment tailored to bring benefits for plant growth and resistance to biotic and abiotic challenges (72). Bioorganic fertilizers promote indigenous soil plant-beneficial consortium to enhance plant disease suppression (73). The design of more efficient biofertilizers to update soil function has important implications for the manipulation of crop microbiomes for sustainable agriculture. Our work provides a comprehensive exploration of microbial interkingdom interactions, hub microbes, and plant genes for the structure of the root microbiome. The results obtained could help design synthetic microbiomes beneficial for plant growth.

## EXPERIMENTAL PROCEDURES

### Root microbiome experiment

Bergelson et al (33, 42) conducted a genome-wide association study (GWAS) for the root microbiome in *Arabidopsis thaliana.* The study included 179 accessions of *A. thaliana,* each measured for the bacterial and fungal abundance of the root microbiota using a 16S/ITS rRNA gene sequencing technique and genotyped for Arabidopsis SNPs by a high-throughput sequencing technology (Table S5).

### Microbial interactions analysis

We chose the 100 most abundant bacterial OTUs and the 100 most abundant fungal OTUs (Table S6) to reconstruct microbial interaction networks using a microbial behavioral network model (45). This model is based on mathematical descriptors of four types of microbe-microbe interactions, mutualism, antagonism, aggression, and altruism, expressed as

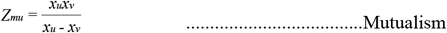

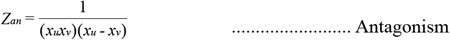

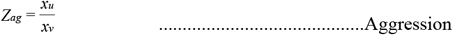

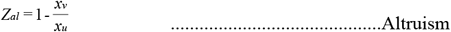

where x_u_ and x_v_ (x_u_ > x_v_, u ≠ v, u, v = 1, ..., m) are the abundance of two microbes u and v and m is the number of microbes. Based on the above equations, we use the corrected microbial abundance to quantify four interactional relationships of two microbes u and v. The descriptor, Z_mu_, can be used to quantify a cooperative relationship (mutualism) between two microbes. The descriptor, Z_an_, can be used to quantify a competitive relationship (antagonism) between two microbes. The descriptor, Z_ag_, can represent the utilization extent (aggression) of a more abundant microbe to a less abundant microbe. The descriptor, Zal, can represent the sacrifice extent (altruism) of a more abundant microbe to a less abundant microbe.

### Microbial networks analysis

By microbial interactions analysis, we can reveal internal workings within the root microbial community. The interaction networks can be visualized using Gephi (https://gephi.org/). We constructed the corresponding network for mutualism, antagonism, aggression, and altruism, respectively. Emergent properties of each network can be calculated in the “igraph” R package (74). We calculated six network indices to describe the features of various networks, including connectivity (Con), closeness (C(u)), betweenness(B(u)), eccentricity (E(u)), eigenvector (G(u)) and Pagerank (P(u)). The specific calculation method was described by Jiang et al. (45). The heat maps of each network index were generated by package *pheatmap* in R (https://CRAN.R-project.org/package=pheatmap). Meanwhile, microbial networks can be used to statistically identify hub taxa. We calculated the degree of each node for every network using the “igraph” R package (74). It is generally believed that hub microbes with a high degree and closeness centrality value play crucial roles in microbial networks (2, 24).

### Mapping microbial network properties

To study how host genes influence root microbiomes, we consider six network property parameters as phenotypic traits that are associated with host SNPs (Single Nucleotide Polymorphisms). We chose those SNPs with MAF > 5%) for association analysis. A regression model of log-transformed phenotypes at a SNP is expressed as

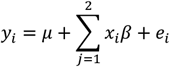

where y_i_ represents the phenotype of the ith host, μ is the mean of the phenotypes over all hosts, x_i_ is the genotype indicator of the ith host which is 0 for high-frequency allele and 1 for low-frequency allele, β is the geneic effect of the SNP and e_i_ is a random error value. Then, we used lm function in R for association analysis from which to get the *P*-value of each SNP. Package *qqman* (https://CRAN.R-project.org/package=qqman) was used to draw the Manhattan plot. By statistical testing, we can find significant SNPs that are associated with each network property.

### QTL networks

Many existing approaches attempt to reveal the genetic architecture of complex traits through identifying key individual genes underlying the traits. However, epistatic interactions among different genes have been increasingly recognized to play an important role in genetic control. Several approaches have been developed to map epistatic interactions based on gene pairwise analysis, failing to systematically chart a network of epistasis involving all genes. More recently, Jiang et al. (46) proposed an analytical procedure of reconstructing epistatic networks from mapping data. This procedure was used to infer QTL networks of the significant SNPs that mediate the emergent properties of microbial networks. At each SNP, we calculated the mean value of each genotype for a network parameter and assigned this value to each *Arabidopsis* accession, transforming the GWAS data structure from its SNP-phenotype illustration to SNP-based genotype representation. We implemented Bayesian networks (BN) to reconstruct genetic networks involving all significant SNPs for each network parameter. The BN-based QTL networks are directed acyclic graphs, encoded by casual SNP-SNP interactions. We identified hub QTLs that play a crucial role in the genetic architecture of plant microbiomes assembly.

## Data availability

The data used can be downloaded at https://doi.org/10.1038/s41598-018-37208-z and the computer code can be freely downloaded at https://github.com/lenahe2006/mSystem for any purpose of research. Also, all questions in data and computation can be addressed to the corresponding author.

## Acknowledgments

We thank Dr. Bergelson and Dr. Horton for supplying their Arabidopsis microbiome and SNP data to us. This work was funded by Natural Science Foundation of China (31971398, 31700633), the Fundamental Research Funds for the Central Universities (2017JC05, 2015ZCQ-SW-06), and Science and Technology Service Network Initiative (KFJ-STS-ZDTP-036).

## Author contributions

X.H. performed data analysis and wrote the manuscript. Q.Z. wrote the R software. L.J. and YJ. critically revised the manuscript. R.W. conceived of the idea and formulated the model. All authors discussed results.

## Conflict of interest

The authors declare that they have no conflict of interest.

## SUPPLEMENTAL MATERIAL

**Table S1 Bacterial and fungal co-occurrence network characteristics and hub microbes.**

**Table S2 The edge information in the four networks.**

**Table S3 Significant QTLs associated with the microbial interaction network.**

**Table S4 The gene enrichment analysis of Hub QTLs.**

**Table S5 The list of accessions.**

**Table S6 The relative abundance of top 100 OTUs in bacteria and fungi.**

**Figure S1** Manhattan plots of GWAS results for *Arabidopsis thaliana* genomic regions associated with emergent properties of root microbial networks. The x-axis showed SNP positions (Mb) and the y-axis was the-log(P-value) resulting from the association test. Each dot in the plot represented an SNP, and a reference line was used on the y-axis to reflect genome-wide significance. The red dotted lines correspond to the chosen significant threshold −log_10_(P)≥ 5. (A) Mutualism, (B) Antagonism, (C) Aggression, and (D) Altruism.

## References

1. Beilsmith K, Thoen M, Brachi B, Gloss A, Khan M, Bergelson J. 2018. Genome-Wide Association studies on the phyllosphere microbiome: embracing complexity in host-microbe interactions. The Plant Journal 97:164–181.

2. Agler MT, Ruhe J, Kroll S, Morhenn C, Kim S-T, Weigel D, Kemen EM. 2016. Microbial Hub Taxa Link Host and Abiotic Factors to Plant Microbiome Variation. PLOS Biology 14:e1002352.

3. Müller D, Vogel C, Bai Y, Vorholt J. 2016. The Plant Microbiota: Systems-Level Insights and Perspectives. Annual review of genetics 50:211–234.

4. Tabrett A, Horton MW. 2020. The influence of host genetics on the microbiome. F1000Research 9:F1000 Faculty Rev-84.

5. Mitter B, Brader G, Pfaffenbichler N, Sessitsch A. 2019. Next generation microbiome applications for crop production — limitations and the need of knowledge-based solutions. Current Opinion in Microbiology 49:59–65.

6. Shelake RM, Pramanik D, Kim J-Y. 2019. Exploration of Plant-Microbe Interactions for Sustainable Agriculture in CRISPR Era. Microorganisms 7:269.

7. Coller E, Cestaro A, Zanzotti R, Bertoldi D, Pindo M, Larger S, Albanese D, Mescalchin E, Donati C. 2019. Microbiome of vineyard soils is shaped by geography and management. Microbiome 7:140.

8. Wagner MR, Lundberg DS, Del Rio TG, Tringe SG, Dangl JL, Mitchell-Olds T. 2016. Host genotype and age shape the leaf and root microbiomes of a wild perennial plant. Nature communications 7:12151–12151.

9. Bakker P, Pieterse C, de Jonge R, Berendsen R. 2018. The Soil-Borne Legacy. Cell 172:1178–1180.

10. de Vries FT, Griffiths RI, Knight CG, Nicolitch O, Williams A. 2020. Harnessing rhizosphere microbiomes for drought-resilient crop production. Science 368:270–274.

11. Harbort CJ, Hashimoto M, Inoue H, Niu Y, Guan R, Rombolà AD, Kopriva S, Voges MJEEE, Sattely ES, Garrido-Oter R, Schulze-Lefert P 2020. Root-Secreted Coumarins and the Microbiota Interact to Improve Iron Nutrition in Arabidopsis. Cell Host & Microbe.

12. Lee S-M, Kong HG, Song G, Ryu C-M. 2020. Disruption of Firmicutes and Actinobacteria abundance in tomato rhizosphere causes the incidence of bacterial wilt disease. The ISME Journal doi:10.1038/s41396-020-00785-x.

13. Tao K, Kelly S, Radutoiu S. 2019. Microbial associations enabling nitrogen acquisition in plants. Current Opinion in Microbiology 49:83–89.

14. Zhang J, Liu Y-X, Zhang N, Hu B, Jin T, Xu H, Qin Y, Yan P, Zhang X, Guo X, Hui J, Cao S, Wang X, Wang C, Wang H, Qu B, Fan G, Yuan L, Garrido-Oter R, Chu C, Bai Y. 2019. NRT1.1B is associated with root microbiota composition and nitrogen use in field-grown rice. Nature Biotechnology 37:676–684.

15. Lu T, Ke M, Lavoie M, Jin Y, Fan X, Zhang Z, Fu Z, Sun L, Gillings M, Penuelas J, Qian H. 2018. Rhizosphere microorganisms can influence the timing of plant flowering. Microbiome 6:231.

16. Panke-Buisse K, Poole AC, Goodrich JK, Ley RE, Kao-Kniffin J. 2015. Selection on soil microbiomes reveals reproducible impacts on plant function. The ISME journal 9:980–989.

17. Mueller E, Wisnoski N, Peralta A, Lennon J. 2019. Microbial rescue effects: How microbiomes can save hosts from extinction. Functional Ecology doi:10.1111/1365-2435.13493.

18. Xu L, Coleman-Derr D. 2019. Causes and consequences of a conserved bacterial root microbiome response to drought stress. Current Opinion in Microbiology 49:1–6.

19. Brachi B, Filiault D, Darme P, Mentec ML, Kerdaffrec E, Rabanal F, Anastasio A, Box M, Duncan S, Morton T, Novikova P, Perisin M, Tsuchimatsu T, Woolley R, Yu M, Dean C, Nordborg M, Holm S, Bergelson J. 2017. Plant genes influence microbial hubs that shape beneficial leaf communities. bioRxiv doi:10.1101/181198:181198.

20. Chen T, Nomura K, Wang X, Sohrabi R, Xu J, Yao L, Paasch BC, Ma L, Kremer J, Cheng Y, Zhang L, Wang N, Wang E, Xin X-F, He SY 2020. A plant genetic network for preventing dysbiosis in the phyllosphere. Nature 580:653–657.

21. Chen Y, Bonkowski M, Shen Y, Griffiths BS, Jiang Y, Wang X, Sun B. 2020. Root ethylene mediates rhizosphere microbial community reconstruction when chemically detecting cyanide produced by neighbouring plants. Microbiome 8:4.

22. Durán P, Thiergart T, Garrido-Oter R, Agler M, Kemen E, Schulze-Lefert P, Hacquard S. 2018. Microbial Interkingdom Interactions in Roots Promote Arabidopsis Survival. Cell 175:973–983.e14.

23. Layeghifard M, Hwang DM, Guttman DS. 2017. Disentangling Interactions in the Microbiome: A Network Perspective. Trends in Microbiology 25:217–228.

24. Xiong C, Zhu Y, Wang J, Singh B, Han L, Shen J, Li P, Wang G, Wu C, Ge A, Zhang L, He J. 2020. Host selection shapes crop microbiome assembly and network complexity. New Phytologist doi:10.1111/nph.16890.

25. Berendsen R, Pieterse C, Bakker P 2012. The rhizosphere microbiome and plant health. Trends in plant science 17:478–86.

26. Pieterse CMJ, de Jonge R, Berendsen RL. 2016. The Soil-Borne Supremacy. Trends in Plant Science 21:171–173.

27. Dahlstrom KM, McRose DL, Newman DK. 2020. Keystone metabolites of crop rhizosphere microbiomes. Current Biology 30:R1131–R1137.

28. Rolfe S, Griffiths J, Ton J. 2019. Crying out for help with root exudates: adaptive mechanisms by which stressed plants assemble health-promoting soil microbiomes. Current Opinion in Microbiology 49:73–82.

29. Thiergart T, Zgadzaj R, Bozsóki Z, Garrido-Oter R, Radutoiu S, Schulze-Lefert P. 2019. Lotus japonicus Symbiosis Genes Impact Microbial Interactions between Symbionts and Multikingdom Commensal Communities. mBio 10:e01833–19.

30. Wagg C, Schlaeppi K, Banerjee S, Kuramae EE, van der Heijden MGA. 2019. Fungal-bacterial diversity and microbiome complexity predict ecosystem functioning. Nature Communications 10:4841.

31. Cordovez V, Dini-Andreote F, Carrion V, Raaijmakers J. 2019. Ecology and Evolution of Plant Microbiomes. Annual Review of Microbiology 73.

32. Getzke F, Thiergart T, Hacquard S. 2019. Contribution of bacterial-fungal balance to plant and animal health. Current Opinion in Microbiology 49:66–72.

33. Bergelson J, Mittelstrass J, Horton M. 2019. Characterizing both bacteria and fungi improves understanding of the Arabidopsis root microbiome. Scientific Reports 9.

34. Trivedi P, Leach JE, Tringe SG, Sa T, Singh BK. 2020. Plant–microbiome interactions: from community assembly to plant health. Nature Reviews Microbiology:1–15.

35. Wang Q, Liu X, Jiang L, Cao Y, Zhan X, Griffin CH, Wu R. 2019. Interrogation of Internal Workings in Microbial Community Assembly: Play a Game through a Behavioral Network? mSystems 4:e00550–19.

36. Fadiji AE, Babalola OO. 2020. Metagenomics methods for the study of plant-associated microbial communities: A review. Journal of Microbiological Methods 170:105860.

37. Banerjee S, Schlaeppi K, van der Heijden MGA. 2018. Keystone taxa as drivers of microbiome structure and functioning. Nature Reviews Microbiology 16:567–576.

38. de Vries FT, Griffiths RI, Bailey M, Craig H, Girlanda M, Gweon HS, Hallin S, Kaisermann A, Keith AM, Kretzschmar M, Lemanceau P, Lumini E, Mason KE, Oliver A, Ostle N, Prosser JI, Thion C, Thomson B, Bardgett RD. 2018. Soil bacterial networks are less stable under drought than fungal networks. Nature Communications 9:3033.

39. Ma B, Wang Y, Ye S, Liu S, Stirling E, Gilbert JA, Faust K, Knight R, Jansson JK, Cardona C, Röttjers L, Xu J. 2020. Earth microbial co-occurrence network reveals interconnection pattern across microbiomes. Microbiome 8:82.

40. Röttjers L, Faust K. 2018. From hairballs to hypotheses - biological insights from microbial networks. FEMS microbiology reviews 42.

41. Kurtz ZD, Müller CL, Miraldi ER, Littman DR, Blaser MJ, Bonneau RA. 2015. Sparse and compositionally robust inference of microbial ecological networks. PLoS Comput Biol 11:e1004226.

42. Horton MW, Bodenhausen N, Beilsmith K, Meng D, Muegge BD, Subramanian S, Vetter MM, Vilhjálmsson BJ, Nordborg M, Gordon JI, Bergelson J. 2014. Genome-wide association study of Arabidopsis thaliana leaf microbial community. Nature communications 5:5320–5320.

43. Jin T, Wang Y, Huang Y, Xu J, Zhang P, Wang N, Liu X, Chu H, Liu G, Jiang H, Li Y, Xu J, Kristiansen K, Xiao L, Zhang Y, Zhang G, Du G, Zhang H, Zou H, Zhang H, Jie Z, Liang S, Jia H, Wan J, Lin D, Li J, Fan G, Yang H, Wang J, Bai Y, Xu X. 2017. Taxonomic structure and functional association of foxtail millet root microbiome. GigaScience 6:1–12.

44. Widder S, Allen RJ, Pfeiffer T, Curtis TP, Wiuf C, Sloan WT, Cordero OX, Brown SP, Momeni B, Shou W, Kettle H, Flint HJ, Haas AF, Laroche B, Kreft J-U, Rainey PB, Freilich S, Schuster S, Milferstedt K, van der Meer JR, Groβkopf T, Huisman J, Free A, Picioreanu C, Quince C, Klapper I, Labarthe S, Smets BF, Wang H, Soyer OS, Isaac Newton Institute F. 2016. Challenges in microbial ecology: building predictive understanding of community function and dynamics. The ISME Journal 10:2557–2568.

45. Jiang L, Liu X, He X, Jin Y, Cao Y, Zhan X, Griffin CH, Gragnoli C, Wu R. 2020. A behavioral model for mapping the genetic architecture of gut-microbiota networks. Gut Microbes doi:10.1080/19490976.2020.1820847:1-15.

46. Jiang L, Xu J, Sang M, Zhang Y, Meixia Y, Zhang H, Wu B, Zhu Y, Xu P, Tai R, Zhao Z, Jiang Y, Dong C, Sun L, Griffin C, Gragnoli C, Wu R. 2019. A Drive to Driven Model of Mapping Intraspecific Interaction Networks. iScience 22.

47. Dai N, Wang W, Patterson SE, Bleecker AB. 2013. The TMK subfamily of receptor-like kinases in Arabidopsis display an essential role in growth and a reduced sensitivity to auxin. PloS one 8:e60990.

48. Strader LC, Culler AH, Cohen JD, Bartel B. 2010. Conversion of Endogenous Indole-3-Butyric Acid to Indole-3-Acetic Acid Drives Cell Expansion in Arabidopsis Seedlings. Plant Physiology 153:1577–1586.

49. Bashandy T, Guilleminot J, Vernoux T, Caparros-Ruiz D, Ljung K, Meyer Y, Reichheld J-P. 2010. Interplay between the NADP-Linked Thioredoxin and Glutathione Systems in *Arabidopsis* Auxin Signaling. The Plant Cell 22:376–391.

50. Zhou H, Zhao J, Cai J, Patil SB. 2017. UBIQUITIN-SPECIFIC PROTEASES function in plant development and stress responses. Plant Molecular Biology 94:565–576.

51. Li L-M, Lü S-Y, Li R-J. 2017. The Arabidopsis endoplasmic reticulum associated degradation pathways are involved in the regulation of heat stress response. Biochemical and Biophysical Research Communications 487.

52. Yamaguchi Y, Huffaker A, Bryan AC, Tax FE, Ryan CA. 2010. PEPR2 Is a Second Receptor for the Pep1 and Pep2 Peptides and Contributes to Defense Responses in Arabidopsis. The Plant Cell 22:508–522.

53. Meyers B, Morgante M, Michelmore R. 2002. TIR-X and TIR-NBS proteins: Two new families related to disease resistance TIR-NBS-LRR proteins encoded in Arabidopsis and other plant genomes. The Plant journal: for cell and molecular biology 32:77–92.

54. Furniss JJ, Grey H, Wang Z, Nomoto M, Jackson L, Tada Y, Spoel SH. 2018. Proteasome-associated HECT-type ubiquitin ligase activity is required for plant immunity. PLOS Pathogens 14:e1007447.

55. Zhang R, Vivanco J, Shen Q. 2017. The unseen rhizosphere root–soil–microbe interactions for crop production. Current Opinion in Microbiology 37:8–14.

56. Ren Y, Xun W, Yan H, Ma A, Xiong W, Shen Q, Zhang R. 2020. Functional compensation dominates the assembly of plant rhizospheric bacterial community. Soil Biology and Biochemistry 150:107968.

57. Martin FM, Uroz S, Barker DG. 2017. Ancestral alliances: Plant mutualistic symbioses with fungi and bacteria. Science 356:eaad4501.

58. Qu Z, Zhao H, Zhang H, Wang Q, Yao Y, Cheng J, Lin Y, Xie J, Fu Y, Jiang D. 2020. Bio-priming with a hypovirulent phytopathogenic fungus enhances the connection and strength of microbial interaction network in rapeseed. npj Biofilms and Microbiomes 6:45.

59. Rodriguez P, Rothballer M, Paul Chowdhury S, Nussbaumer T, Gutjahr C, Falter-Braun P 2019. Systems Biology of Plant-Microbiome Interactions. Molecular Plant 12.

60. Yuanyuan X, Chen H, Yang J, Liu M, Huang B. 2018. Distinct patterns and processes of abundant and rare eukaryotic plankton communities following a reservoir cyanobacterial bloom. The ISME Journal 12.

61. Deng Y, Jiang Y-H, Yang Y, He Z, Luo F, Zhou J. 2012. Molecular ecological network analyses. BMC Bioinformatics 13:113.

62. Pratama AA, Terpstra J, de Oliveria ALM, Salles JF. 2020. The Role of Rhizosphere Bacteriophages in Plant Health. Trends in Microbiology 28:709–718.

63. Xiong W, Song Y, Yang K, gu Y, Wei Z, Kowalchuk G, Xu Y, Jousset A, Shen Q, Geisen S. 2020. Rhizosphere protists are key determinants of plant health. Microbiome 8.

64. Shi Y, Delgado-Baquerizo M, Li Y, Yang Y, Penuelas J, Chu H. 2020. Abundance of kinless hubs within soil microbial networks are associated with high functional potential in agricultural ecosystems. Environment International 142:105869.

65. Liu H, Brettell LE, Singh B. 2020. Linking the Phyllosphere Microbiome to Plant Health. Trends in Plant Science 25:841–844.

66. Lawson C, Harcombe W, Hatzenpichler R, Lindemann S, Löffler F, O’Malley M, Martín H, Pfleger B, Raskin L, Venturelli O, Weissbrodt D, Noguera D, McMahon K. 2019. Common principles and best practices for engineering microbiomes. Nature Reviews Microbiology 17:1–17.

67. Bai Y, Müller D, Srinivas G, Garrido-Oter R, Potthoff E, Rott M, Dombrowski N, Münch P, Spaepen S, Remus-Emsermann M, Huettel B, McHardy A, Vorholt J, Schulze-Lefert P 2015. Functional overlap of the Arabidopsis leaf and root microbiota. Nature 528.

68. Niu B, Paulson JN, Zheng X, Kolter R. 2017. Simplified and representative bacterial community of maize roots. Proceedings of the National Academy of Sciences 114:E2450–E2459.

69. Lebeis SL, Paredes SH, Lundberg DS, Breakfield N, Gehring J, Mcdonald M, Malfatti S, Del Rio TG, Jones CD, Tringe SG. 2015. Salicylic acid modulates colonization of the root microbiome by specific bacterial taxa. Science 349:860–4.

70. Liu Y-X, Qin Y, Bai Y. 2019. Reductionist synthetic community approaches in root microbiome research. Current Opinion in Microbiology 49:97–102.

71. Fitzpatrick CR, Salas-González I, Conway JM, Finkel OM, Gilbert S, Russ D, Teixeira PJPL, Dangl JL. 2020. The Plant Microbiome: From Ecology to Reductionism and Beyond. Annual Review of Microbiology 74:81–100.

72. Lareen A, Burton F, Schäfer P 2016. Plant root-microbe communication in shaping root microbiomes. Plant Molecular Biology 90.

73. Tao C, Li R, Xiong W, Shen Z, Liu S, Wang B, Ruan Y, Geisen S, Shen Q, Kowalchuk GA. 2020. Bio-organic fertilizers stimulate indigenous soil Pseudomonas populations to enhance plant disease suppression. Microbiome 8:137.

74. Csardi G, Nepusz T 2005. The Igraph Software Package for Complex Network Research. InterJournal Complex Systems:1695.

